# Mismatch repair deficiency predicts response to HER2 blockade in HER2-negative breast cancer

**DOI:** 10.1101/2020.06.08.128090

**Authors:** Nindo B Punturi, Sinem Seker, Vaishnavi Devarakonda, Rashi Kalra, Ching-Hui Chen, Aloran Mazumder, Shunqiang Li, Tina Primeau, Matthew J Ellis, Shyam M Kavuri, Svasti Haricharan

## Abstract

Estrogen receptor positive (ER^+^) breast cancer is a leading cause of cancer-related death globally. Resistance to standard of care endocrine treatment occurs in at least 30% of ER^+^ breast cancer patients resulting in ~40,000 deaths every year in the US alone. Preclinical studies strongly implicate activation of growth factor receptor, HER2 in endocrine treatment resistance of ER^+^ breast cancer that is HER2^-^ at diagnosis^1,2^. However, clinical trials of pan-HER inhibitors in ER^+^/HER2^-^ patients have disappointed, likely due to a lack of predictive biomarkers^3-6^. Here we demonstrate that loss of *MLH1*, a principal mismatch repair gene, causally activates HER2 in ER^+^/HER2^-^ breast cancer upon endocrine treatment. Additionally, we show that HER2 activation is indispensable for endocrine treatment resistant growth of MLH1^-^ cells *in vitro* and *in vivo.* Consequently, inhibiting HER2 restores sensitivity to endocrine treatment in multiple experimental models including patient-derived xenograft tumors. Patient data from multiple clinical datasets (TCGA, METABRIC, Alliance (Z1031) and E-GEOD-28826) supports an association between *MLH1* loss, HER2 upregulation, and sensitivity to trastuzumab in endocrine treatment-resistant ER^+^/HER2^-^ patients. These results provide strong rationale that MLH1 could serve as a first-in-class predictive marker of sensitivity to combinatorial treatment with endocrine drugs and HER inhibitors in endocrine treatment-resistant ER^+^/HER2^-^ breast cancer patients. Implications of this study extend beyond breast cancer to Lynch Syndrome cancers.

**One Sentence Summary:** Defective mismatch repair activates HER2 in HER2-negative breast cancer cells and renders them susceptible to HER2 inhibitors.

## Introduction

Estrogen receptor (ER) positive breast cancer is one of the most common cancers in women worldwide^7^. ER^+^ breast cancer patients are treated with endocrine therapy which interrupts ER signaling^8^. A subset of ER^+^ breast tumors also amplify the tyrosine kinase receptor and oncogene, HER2^1,2^ These ER^+^/HER^+^ breast cancer patients are less responsive to endocrine therapy but respond extremely well to combinatorial treatment with a HER2 inhibitor, a game changing discovery^9^. However, the majority of ER^+^ breast cancer is HER2^-^ at diagnosis, and while ~70% of ER^+^/HER2^-^ breast cancer patients respond well to endocrine treatment, ~30% of patients become resistant to endocrine treatment resulting in relapse, metastasis and death^8,10^.

The discovery that HER2 amplification induces endocrine therapy resistance in ER^+^ breast cancer spurred research into other means of HER2 activation. These studies identified *HER2* mutation and phosphorylation as mechanisms by which ER^+^ HER2 non-amplified (henceforth referred to as ER^+^ HER2^-^) breast cancer cells could resist endocrine treatment^2,11^. However, translation of these findings proved challenging with results from clinical trials not living up to preclinical promise^3,6^. There is recognition now that this is likely because only a subset of ER^+^ breast cancers activate HER2 to resist endocrine therapy. Finding this subset is further complicated by the fact that ER^+^/HER2^-^ breast cancer cells likely activate HER2 only in response to endocrine treatment, making identification of these patient cohorts before administering treatment challenging. Without a means of identifying this patient subset, it is difficult to design a clinical trial with sufficient resolution to uncover real improvement in patient outcome.

Continuing efforts to identify alternative therapies for endocrine treatment resistance have resulted in few improvements in the clinic. The only targeted therapy to prove effective to date is the recently discovered group of CDK4/6 inhibitors^12^. However, these inhibitors have to be administered constantly to be effective, and are, therefore, a financially and physically costly treatment modality that postpones resistance, metastasis and death but does not remove this threat^13^. Moreover, some endocrine treatment resistant patients do not respond to CDK4/6 inhibitors^14^. Hope of curing endocrine treatment resistant patients with HER2 inhibitors, therefore, remains a tantalizing challenge with immense clinical impact.

Defects in the MutL complex of mismatch repair, comprised of *MLH1* and *PMS2*, were recently identified as drivers of endocrine treatment resistance in 15-17% of ER^+^/HER2^-^ breast cancer patients^15,16^. Mismatch repair is a fundamental DNA repair pathway conserved between pro- and eukaryotes, and essential for guarding the genome during cellular replication^17^. Here, we demonstrate a novel role for MutL loss in activating HER2 in ER^+^ HER2^-^ cells when exposed to endocrine treatments. Moreover, using multiple experimental model systems we provide strong evidence for MutL loss as a stratifier for response to HER inhibitors in endocrine treatment resistant, nominally HER2^-^ ER^+^ breast cancer patients.

## Results

### Loss of mismatch repair associates with HER2 activation in HER2^-^ breast cancer cells

To understand mechanisms underlying MutL loss-induced endocrine treatment resistance, we conducted a proteomics screen. We used reverse phase protein array to compare ER^+^/HER2^-^ MCF7 breast cancer cells engineered to carry shRNA against *MLH1* or *PMS2* against control isogenic cells with shRNA against Luciferase. All cell lines were treated with and without endocrine treatment in the form of fulvestrant, an estrogen receptor down-regulator to understand the impact of ER loss in this system. We identified significant upregulation of phosphorylated HER2 (pHER2) in sh*MLH1* and sh*PMS2* MCF7 cells but not in sh*Luc* cells upon fulvestrant treatment (Fig S1A). To test whether such an association between MutL loss and HER2 activation could be detected in patient tumors, we analyzed ER^+^ breast tumors that were nominally HER2^-^ (non-amplified) from two independent datasets: METABRIC and TCGA. In both cases, we observed that ~25% of MutL^-^ patient tumors had relatively high RNA levels of HER2 compared to ~10% of MutL^+^ patient tumors (Fig 1A, contextualized in Fig S1B-C). MutL^-^ patient tumors with relatively high *HER2* RNA also associated with significantly worse disease-specific survival in METABRIC (Fig 1B) and in TCGA (Fig S1D). MutL loss as assayed by low gene expression levels is not an artifact of low basal proliferation rate, since RNA levels of *MKI67* (a proliferation marker) are comparable between patient tumors designated as MutL^-^ and MutL^+^ (Figs S1E-F). Upregulation of HER2 in MutL^-^ patient tumors also independently prognosticated worse disease-specific survival in Cox Proportional Hazards analyses (Fig 1C). These data suggest that the association between MutL loss and HER2 upregulation is of clinical relevance.

**Figure 1:**
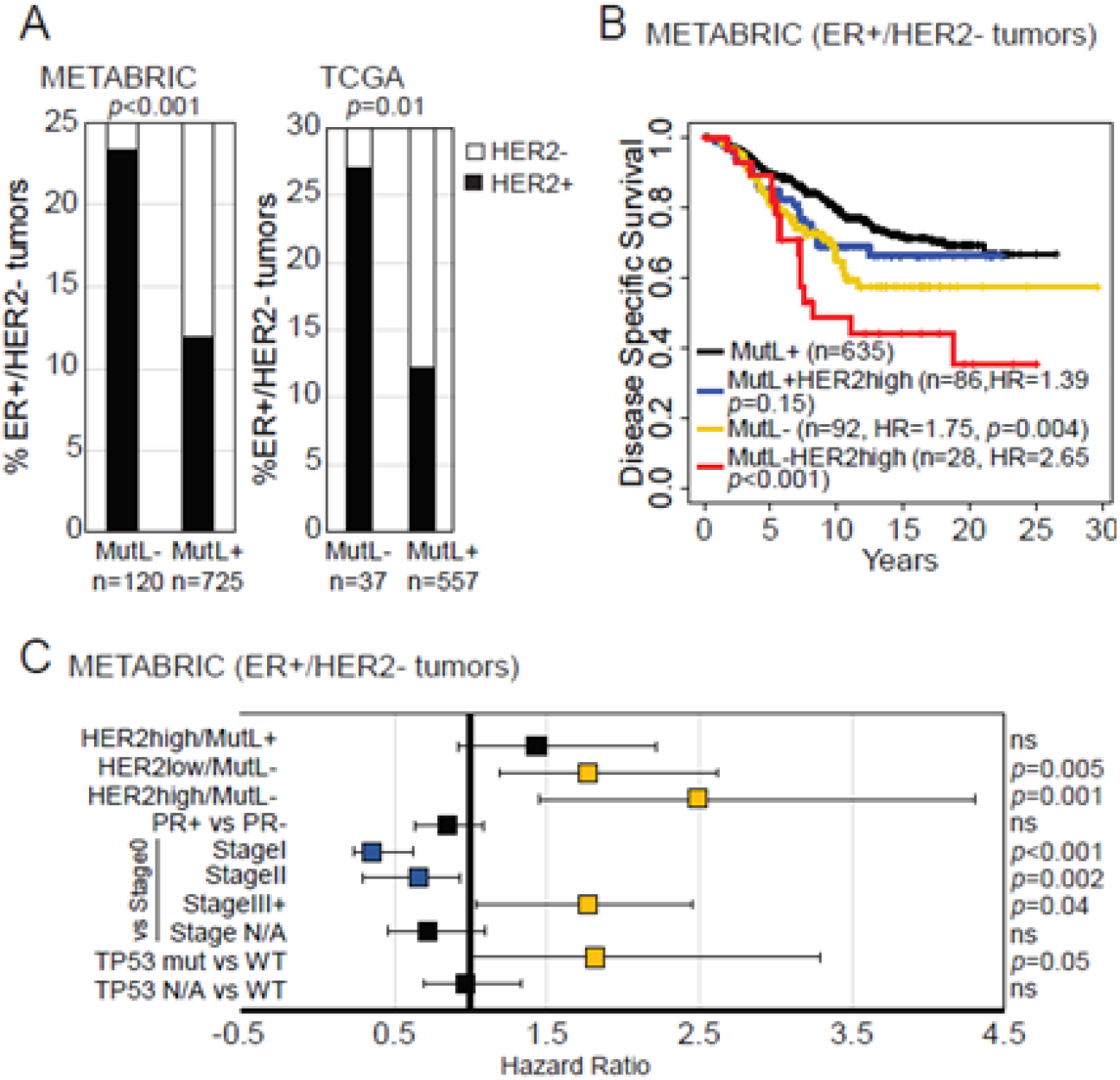
ER^+^, nominally HER2^-^ (non-amplified) breast cancer patients whose tumors are MutL^-^ have relatively high RNA levels of *HER2* and associate with significantly worse disease-specific survival. (A) Incidence of tumors with high *HER2* RNA levels within MutL^-^ and MutL^+^ ER^+^/HER2^-^ breast tumors from METABRIC and TCGA. Pearson Chi-Square test identified p-values. Contextualization with HER2^+^ subset in Fig S1B-C. (B-C) Kaplan-Meier survival curves (B) and proportional hazard assessment (C) demonstrating differences in disease-specific survival between specified groups within the ER^+^/nominally HER2^-^ breast tumor cohort from METABRIC. Cox Regression analysis identified p-values. Supporting data from TCGA presented in Fig S1D and proliferation controls in Figs S1E-F.

### Inhibition of mismatch repair induces activation of HER2 in ER^+^/HER2^-^ breast cancer cells in response to endocrine treatment

We next tested the causality of this relationship in two independent cell line models of ER^+^/HER2^-^ breast cancer: MCF7 and T47D. In both cell lines, Western blotting identified higher levels of pHER2 in sh*MLH1* isogenic lines relative to sh*Luc* at baseline with further increase upon treatment with ER degrader, fulvestrant (Fig 2A, Fig S2A). Downstream signaling to pAkt and pS6k was also upregulated in sh*MLH1* cells after fulvestrant treatment (Fig 2A). Additionally, we confirmed increased pHER2 at the membrane of sh*MLH1* cells after fulvestrant treatment using both immunofluorescence (Fig 2B) and flow cytometry (Fig S2B). Increase in membrane HER2 in sh*MLH1* cells after exposure to endocrine treatment was even more striking in xenograft tumors from MCF7 sh*Luc* and sh*MLH1* cells (Fig 2C). Importantly, the same increase in membrane HER2 levels after fulvestrant treatment was seen in tumors from an ER^+^/HER2^-^ patient-derived xenograft (PDX) model of MutL loss (WHIM20^15,18^) (Fig 2D). These data indicate a causal link between MutL loss and HER2 activation catalyzed by endocrine treatment.

**Figure 2:**
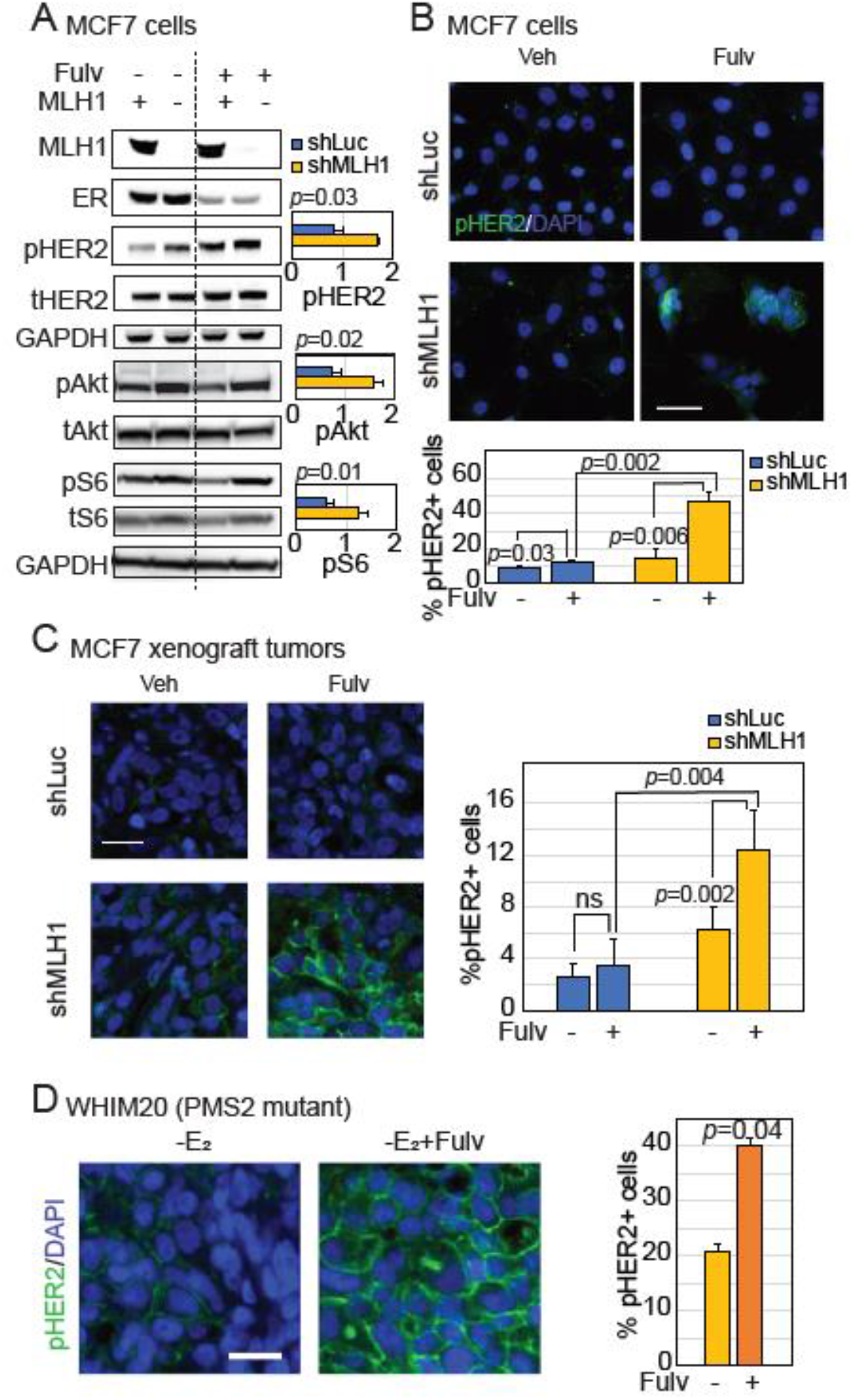
MLH1 loss in ER^+^, nominally HER2^-^ breast cancer cells causally increases membrane-bound HER2. (A) Western blots demonstrating increases in pHER2 and associated downstream signaling in sh*MLH1* MCF7 cells treated with fulvestrant relative to sh*Luc* cells. Quantification of four independent replicates conducted through ImageJ in accompanying bar graphs. Validation in T47D cells in Fig S2A. (B-D) Immunofluorescent staining for pHER2 in MCF7 sh*Luc* and sh*MLH1* cells *in vitro* (B), MCF7 sh*Luc* and sh*MLH1* xenograft tumors (C), and in WHIM20, *PMS2* mutant, ER^+^/HER2^-^ PDX tumors (D), grown with or without fulvestrant. Accompanying quantification presented as bar graphs. All columns indicate the mean and error bars the standard deviation. At least three independent experiments or five independent tumors from each group were quantified. Student’s t-test determined p-values. Supporting data from FACS analysis is presented in Fig S2B. Scale bars represent 50μ.

Since MutL^-^ ER^+^ tumors have a higher mutation load than MutL^+^ tumors^15,19^ we tested whether *HER2* activation in these tumors occurred *via* activating mutations^11^. However, frequency of *HER2* mutations did not significantly associate with MutL status, if anything appearing lower in MutL^-^ ER^+^ tumors than in MutL^+^ ones (Fig S2C). This suggests that MutL loss induces HER2 activation through other non-mutational mechanisms.

### HER2 activation is required for endocrine treatment resistant growth of MutL^-^ ER^+^/HER2^-^ breast cancer cells

To test this link genetically, we used siRNA to decrease endogenous *HER2* in MCF7 sh*Luc* and sh*MLH1* cells, and then assayed growth in presence of fulvestrant. We observed complete rescue of endocrine treatment sensitivity in sh*MLH1* cells transfected with si*HER2*, with no observable change in endocrine treatment response in sh*Luc* cells under the same conditions (Fig 3A-B). This dependence on HER2 for endocrine treatment resistant growth in sh*MLH1* cells was also observed with two additional endocrine therapies: tamoxifen, an ER modulator (Fig S3A), and estrogen deprivation (Fig S3B), a surrogate for aromatase inhibitors used in clinic. These data indicate that HER2 activation is necessary for endocrine treatment resistance observed in MutL^-^ ER^+^/HER2^-^ breast cancer cells.

**Figure 3:**
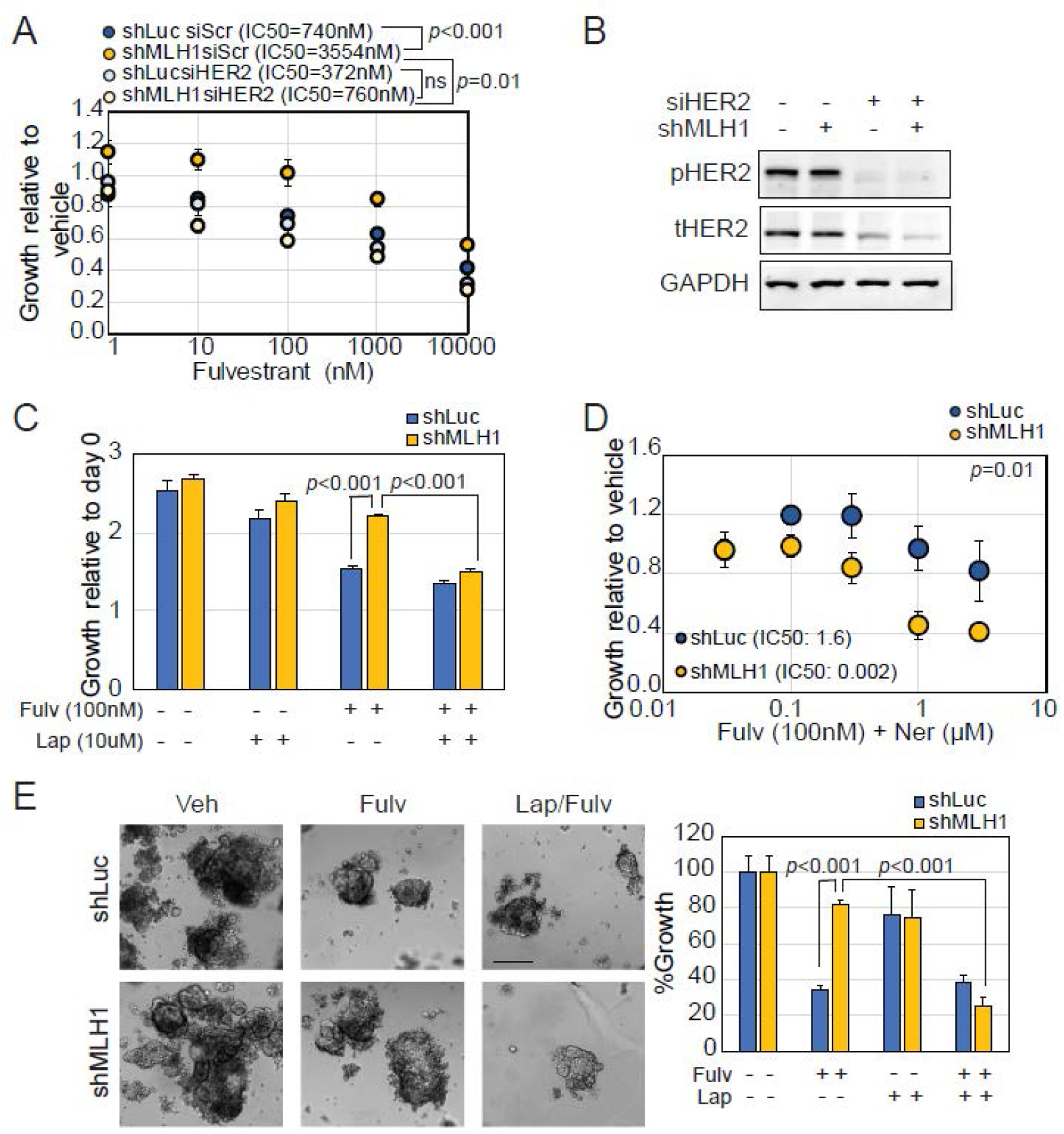
HER2 is required for endocrine treatment resistant growth of ER^+^ MLH1^-^ breast cancer cells. (A-B) Knockdown of endogenous *HER2* using siRNA against *HER2* or a scrambled control in MCF7 sh*Luc* and sh*MLH1* cells validated by Western blotting (B) and followed by 2D growth assays for dose response to fulvestrant treatment (A). Supporting data demonstrating similar response to tamoxifen and estrogen deprivation in Fig S3A-B. (C) Growth of MCF7 sh*Luc* and sh*MLH1* cells in response to specified therapeutic combinations represented as a bar graph. Supporting data from T47D in Fig S3C. (D) Dose curve demonstrating response to neratinib and fulvestrant in MCF7 sh*Luc* and sh*MLH1* cells. Supporting data demonstrating similar results in T47D cells and in response to tamoxifen in Fig S3D-E. (E) 3D growth in Matrigel of MCF7 sh*Luc* and sh*MLH1* cells in response to specified treatments. Representative images shown alongside quantification. Supporting data in T47D cells in Fig S3F. For dose curve experiment, IC50 values were determined over three independent experiments and compared for statistical differences. Circles represent mean growth relative to vehicle treated cells over 7 days of treatment and error bars the standard deviation. For bar graphs, columns represent the mean and error bars the standard deviation. All statistical comparisons used Student’s two-tailed t-test. All experiments were conducted >2 times.

In keeping with this observation, both MCF7 (Fig 3C) and T47D (Fig S3C) sh*MLH1* cells grown in 2D were sensitive to combinatorial administration of fulvestrant and lapatinib, a HER inhibitor used in clinic. Additionally, MCF7 (Fig 3D) and T47D (Fig S3D) sh*MLH1* cells also demonstrated increased sensitivity to fulvestrant when treated with neratinib, another HER inhibitor. Similar results were obtained when neratinib was combined with tamoxifen treatment (Fig S3E). Finally, both MCF7 (Fig 3E) and T47D (Fig S3F) sh*MLH1* cells demonstrated persistent 3D growth relative to sh*Luc* cells in response to fulvestrant, but this growth was significantly suppressed by adding lapatinib. These data suggest that loss of MutL predisposes ER^+^/HER2^-^ breast cancer cells to respond to HER2 inhibitors in concert with endocrine therapies.

To test this proposition *in vivo*, we randomized mice with MCF7 sh*MLH1* xenograft tumors into four treatment arms: control, fulvestrant, lapatinib, and a combination of fulvestrant and lapatinib. All arms were deprived of estrogen supplementation at randomization. As expected from previous experiments^15^, we observed estrogen independent and fulvestrant resistant growth in MCF7 sh*MLH1* tumors, and little response to lapatinib alone (Fig 4A). However, there was striking response with tumor shrinkage to the combination of fulvestrant and lapatinib (Fig 4A). Congruently, assessment of Ki67 in resultant tumors demonstrated direct correlation between proliferation and HER2 positivity in tumors treated with fulvestrant, but neither in the control group nor in the combination group treated with lapatinib and fulvestrant (Fig S4A).

**Figure 4:**
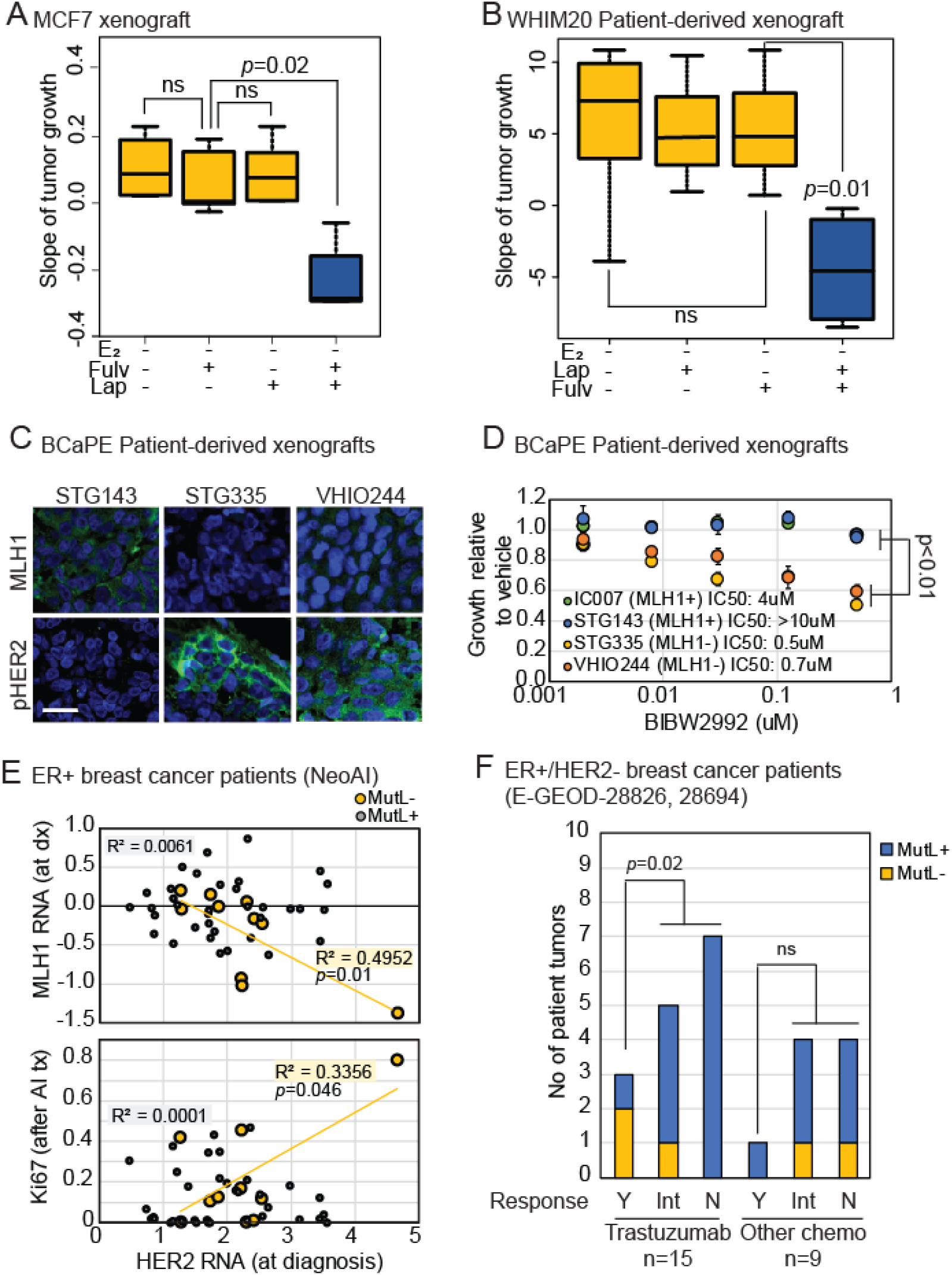
MLH1 loss predicts sensitivity to HER2 inhibitors in endocrine treatment resistant ER^+^, nominally HER2^-^ breast cancer cells *in vivo* and in patient tumors. (A-B) *In vivo* xenograft experiments of MCF7 sh*MLH1* cells (A) and WHIM20 PDX line (B) demonstrating response in tumor growth to specified treatments. Box plots quantify slope of tumor growth with the lines representing the mean and error bars the standard deviation. More than 5 mice were included in each treatment group. Student’s t-test determined p-values. Supporting data in Fig S4A-C. (C-D) Immunofluorescence depicting low protein levels of MLH1 in two ER^+^/HER2^-^ PDX lines associating with increased HER2 activation after estrogen deprivation (C) and increased sensitivity to HER2 inhibition when grown ex vivo (D). Scale bar=50μ. IC50 calculated using regression analysis over three independent experiments and compared using Student’s T-test. (E) Regression analysis demonstrating inverse correlation between *MLH1* RNA levels (top)/Post-endocrine treatment Ki67 (bottom) with *HER2* RNA levels in ER^+^/HER2^-^ patient tumors from Z1031. Supporting data in Fig S4D. (F) Categorical analysis supporting increased sensitivity of advanced ER^+^/MutL^-^ patient tumors to trastuzumab given in combination with other chemotherapy. Supporting data in Fig S4E. Y, Yes; Int, Intermediate; N, No. Fisher’s Exact test determined p-values. For all regression analyses, individual p-values and multiple adjusted R^2^ values derived using a linear model.

### Loss of mismatch repair increases sensitivity to HER2 inhibitors in vivo and in patient tumors

We next tested whether MutL defects had similar associations with sensitivity to HER2 inhibitors in PDX tumors. *In vivo* growth of WHIM20, *PMS2* mutant, ER^+^ PDX tumors xenografted into mouse mammary fat pads demonstrated a similar pattern of tumor regression in response to combination of lapatinib and fulvestrant but not in response to either treatment alone (Fig 4B).

Additionally, two ER^+^/HER2^-^ PDX lines^20^ with loss of nuclear MLH1 also had increased HER2 activation (Fig 4C) and increased sensitivity to BIBW2992 or afatinib, a second generation pan HER inhibitor, as assayed by *ex vivo* 3D growth (Fig 4D). We also observed significant correlation between sensitivity to three HER inhibitors, including lapatinib, and low RNA levels of*MLH1/PMS2* across seven PDX models of luminal breast cancer^20^ grown in estrogen deprived conditions (Fig S4B). Of note, there was no such correlation across 11 PDX models of basal-like breast cancer (Fig S4C). Together, these data demonstrate that MutL loss predisposes ER^+^/HER2^-^ breast tumors to respond to a combination of HER inhibitors and endocrine treatment in three independent PDX lines.

We validated our findings in patient tumors from Z1031^21^, which consists of ‘omics data from ER^+^/HER2^-^ breast cancer patients whose tumors were biopsied at diagnosis and after 4-6 weeks of neoadjuvant aromatase inhibitor treatment. We first confirmed inverse association between RNA levels of *HER2* and *MLH1* in these tumors at diagnosis (Fig 4E), indicating that tumors with low *MLH1* were likely to have relatively higher *HER2* at baseline (as observed in our experimental model systems). Next, we identified direct association between RNA levels of *HER2* and proliferation as measured by immunohistochemistry for Ki67 after endocrine treatment (Fig 4E). Importantly, this association was restricted to MutL^-^ tumors (Fig 4E). These data suggest that loss of MutL induces HER2-associated proliferation in ER^+^/HER2^-^ breast cancer cells treated with endocrine intervention. As an additional control, we found no significant associations between levels of *HER2* RNA those of another mismatch repair gene, *MSH2*, which is not part of the MutL complex (Fig S4D). This specificity increases confidence in the association between HER2 activation and MutL loss. Secondly, association between *HER2* RNA levels and Ki67 in MutL^-^ ER^+^/HER2^-^ breast tumors is only significant after exposure to endocrine treatment and not in pre-treatment biopsies (Fig S4D). Again, we confirmed that loss of MutL in patient tumors was not merely a consequence of low proliferation (Fig S4E). This attests to the role of endocrine therapy in catalyzing reliance on HER2 for proliferation in these tumors.

We also analyzed a second dataset^22^ where metastatic, treatment-resistant breast cancer patients, irrespective of HER2 status, were randomized to two arms of treatment: anthracyclines and taxanes or anthracyclines, taxanes and trastuzumab, a HER2 inhibitor. From this dataset, we parsed the subset of patients whose cancer was ER^+^/HER2^-^ for further analysis. Strikingly, all patients with MutL^-^ ER^+^/HER2^-^ breast cancer demonstrated at least partial response to trastuzumab, compared to less than half of patients with MutL^+^ ER^+^/HER2^-^ disease (Fig 4F). Additionally, 2/3^rd^ of MutL^-^ patients had complete response to the trastuzumab combination compared to less than a tenth of MutL^+^ patients (Fig 4F). This disparate response was only observed in the treatment group where trastuzumab was added to the chemotherapy administered to patients. Concomitant downregulation of *HER2* RNA in response to the trastuzumab combination, but not in response to anthracyclines/taxanes alone was confirmed in the MutL^-^ ER^+^/HER2^-^ tumors (Fig S4F). These data, while of small sample size, provide preliminary support for a role for MutL loss in sensitizing endocrine treatment-resistant ER^+^/HER2^-^ breast cancer to a combination of HER2 inhibitors and endocrine treatment.

## Discussion

Taken together, results presented here suggest that *MLH1*/*PMS2* downregulation could constitute an exciting, new predictive marker for response to HER2 inhibition in endocrine treatment-resistant ER^+^/HER2^-^ breast cancer. The only other biomarkers proposed to predict response to HER2 inhibitors in the endocrine treatment resistant ER^+^/HER2^-^ setting is low ER/PR but these markers have mixed associations across clinical trials decreasing their feasibility for clinical use^3,23^. The impact of the discovery presented here could be substantial, given recent publications suggesting that protein level loss of MLH1 and PMS2 occurs in 15-17% of ER^+^/HER2^-^ breast cancer^24^. Importantly, it is clinically feasible to assess this marker at diagnosis and use it to tailor therapy since diagnostic assays for MLH1 and PMS2 loss are routinely implemented in clinic for colorectal and endometrial cancer patients^25,26^. Coopting these diagnostic tests for ER^+^/HER2^-^ breast cancer patients could benefit a significant subset of patients in a relative short period of time.

The mechanism underlying HER2 activation in response to endocrine treatment in MutL^-^ ER^+^ breast cancer cells remains elusive. Lack of any association between *HER2* mutations and MutL loss is striking and suggests the existence of alternative mechanisms of HER2 activation. Data from Western blots suggest that total HER2 levels do not alter significantly with MutL loss, thereby supporting the existence of a post-translational mechanism. However, undeniably, baseline levels of HER2 appear higher in ER^+^/HER2^-^ MutL^-^ breast cancer cells in patient tumor gene expression data, although many orders lower than levels in HER2-amplified patient tumors. Also, both immunofluorescence and Western blotting of MutL-cell lines and PDXs indicates higher HER2 protein levels at baseline, suggesting a more complex mechanistic underpinning. The link between loss of a DNA damage repair pathway and growth factor receptor activation is certainly intriguing and requires further investigation.

As is often the case in translational research, a significant limitation of this study is the lack of specific clinical trial data in which to test the hypothesis raised by the molecular biology described above. Very few clinical trials have been performed to test efficacy of HER inhibitors in ER^+^/HER2^-^ breast cancer patients^3,4^. None of these trials include transcriptomic or genomic data accrual from tumor biopsies and since MutL status is not routinely tested in clinic for breast cancer patients, this data is missing from all existing trials. Due to failure of these trials to demonstrate positive clinical impact, no further trials were pursued. The strength of preclinical data presented here and the strong associations observed in clinical trial data, albeit limited by sample size, provide a compelling case for revisiting HER2 inhibitors in the ER^+^/HER2^-^ breast cancer setting but this time in context of MutL status.

These results also have significance beyond ER^+^ breast cancer. Our data provide support for a recent report on Lynch syndrome colorectal cancer suggesting a link between loss of mismatch repair and response to HER inhibitors^27^. Lynch syndrome is one of the most common causes of inherited cancers at many sites and is caused by hereditary defects in mismatch repair genes^28^. In addition, mismatch repair loss drives a significant proportion of sporadic colorectal, ovarian, and endometrial cancer^29^. If *MLH1/PMS2* loss serves as a predictive marker for sensitivity to HER2 inhibitors across cancer types, the already routine identification of these markers in these other cancer types can be married to a clinically feasible targeted therapy.

## Materials and Methods

### Cell Lines, Mice, CRISPR, si/shRNA Transfection, and Growth Assays

Cell lines were obtained from the ATCC (2015) and maintained and validated as previously reported^30^. *Mycoplasma* tests were performed on parent cell lines and stable cell lines every 6 months (latest test: 02/19) with the Lonza Mycoalert Plus Kit (cat# LT07-710) as per the manufacturer’s instructions. Cell lines were discarded at <25 passages, and fresh vials were thawed out. Key experiments were repeated with each fresh thaw. Transient transfection with siRNA against HER2 was conducted as previously^30^, and siRNA pools were purchased from Sigma-Aldrich. Stable cell lines were maintained in presence of specified antibiotics at recommended concentrations. Growth assays were conducted in triplicate and repeated independently as previously using Alamar blue to identify cell viability^15^. Growth assay results were plotted as fold change in growth from days 1 to 7 and normalized as specified. Threedimensional growth assays were conducted over 4 to 6 weeks with weekly drug treatments as described previously^11^. Images were captured when colonies had established (at 2 weeks), and then treatment was administered, with images taken again at 1 and 3 weeks post treatment. Fold change in area of colonies was calculated over time and represented as %growth. Tumor growth assays *in vivo* were carried out as described previously^15^ by injecting 2 to 5 × 10^6^ MCF7 cells into the L4 mammary fat pad/mouse. Mice for the MCF7 experiments were 4- to 6-week athymic nu/nu female mice (Envigo). For WHIM20 PDX experiments, 6- to 8-week female SCID/Bg mice were purchased from Jackson laboratory. Tumor volume was measured twice or thrice weekly using calipers to make 2 diametric measurements. Tumors were randomized for treatment at 50 to 150 mm^3^ volume for MCF7 xenografts and 100-300 mm^3^ volume for WHIM20 PDX experiments. Tumors were harvested at <2 cm diameter and were embedded in paraffin blocks, OCT, and snap-frozen as described previously^31^. Mice that died within 3 weeks of tumor growth rate experiments were excluded from analysis. For all mouse experiments, investigator was blinded to groups and to outcomes. STG335, STG143 and VHIO244 PDX experiment results were kindly provided by the BCaPE consortium, but tumor sections were stained in house. All mouse experiments were performed according to the Institutional Animal Care and Use Committee rules and regulations (protocol# AN-6934).

### Inhibitors and Agonists

All drugs were maintained as stock solutions in DMSO, and stock solutions were stored at −80 and working stocks at −20 unless otherwise mentioned. 4-OHT (Sigma-Aldrich, cat# H7904) and fulvestrant (SelleckChem, cat# I4409) were purchased, and stocks were diluted to 10 mmol/L working stocks for all experiments other than dose curves, where specified concentrations were used. For all experiments, cells were treated 24 hours after plating, and thereafter every 48 hours until completion of experiment. For mouse xenograft experiments, fulvestrant concentrations of 250 mg/kg body weight were prepared in corn oil, freshly on day of injection and administered subcutaneously. Beta-estradiol was purchased from Sigma-Aldrich (cat# E8875), maintained in sterile, nuclease-free water, and diluted to obtain 10 mmol/L stocks for *in vitro* experiments. For mouse xenograft experiments, 17 β-estradiol was maintained in 200-proof ethanol at 2.7 mg/ml stock solution and added to drinking water twice a week at a final concentration of 8 μg/mL (cat# E2758; Sigma). Lapatinib (SelleckChem, cat#S2111) and Neratinib were used at specified concentrations. Lapatinib tablets were used at 100 mg/kg in chow from Research Diets, Inc for tumor growth assays.

### Flow cytometry, Immunostaining and Microscopy

Flow cytometry for membrane bound HER2 was performed based on manufacturer’s instructions. After fulvestrant treatment, cells were detached from plates using StemPro™ Accutase™ Cell Dissociation Reagent (cat#A1110501). Cells were washed with chilled PBS and suspended in antibody solution, as per the manufacturer’s instructions, in 5 mL flow cytometry tubes and incubated on ice for 20 minutes. Live cells were then run through BD Accuri to assess only membrane bound HER2 protein levels. IF was performed based on the manufacturer’s instructions. Cells were washed in PBS; fixed for 20 minutes at room temperature in 4% PFA; blocked for 1 hour at room temperature in 5% goat serum and 1% Triton X-100 in 1x PBS; incubated with primary antibody overnight at 4 degrees in 1% goat serum and 1% Triton X-100 in 1x PBS antibody diluent; incubated with secondary antibody in diluent for 1 hour at RT; and then mounted with DAPI-containing mounting media (cat# P36935). Tumor section staining was done as before. Briefly, slides were incubated at 65degrees for 4 hours and deparaffinized. Antigen retrieval was done with 10mM Sodium Citrate (pH 6) for 25 minutes in pressure cooker. Hereafter the cells were treated the same as the 2D IF. Primary antibodies used include pHER2 (EMD millipore; cat# 06-229; 1:200) and Ki67 (Novus Biologics, cat# NB500-170SS, 1:250). Cells were treated with fulvestrant for 24 hours before evaluation. Fluorescent images were captured with a Nikon microscope and quantified with ImageJ. Representative images were translated into figures using Adobe Photoshop and Adobe Illustrator.

### Protein Analyses

Western blotting was conducted as previously described^30^. Cells were exposed to 18-24 hours of fulvestrant treatment administered 40 hours after plating. For pHER2 Western blots, primary antibody was incubated for 48 hours at 4degrees. For all other antibodies, primary incubation was 2 hours at room temperature. All antibodies diluted in 1x TBST and incubated overnight at 4°C. Antibodies used were pHER2 Y1196 (D66B7) (Cell Signaling; cat# 6942S), total HER2 (Thermo Scientific; NeoMarkers; cat# MS-730-P1ABX), pAkt S473 (D9E) XP (Cell Signaling; cat#4060S), total Akt (Cell Signaling; cat#9272S), pS6 (S235/236) (Cell Signaling; cat# 2211S), total S6 (5G10) (Cell Signaling; cat# 2217S), MLH1 (1:2,000, Sigma-Aldrich; cat# WH0004292M2), ER clone 60C (EMD Millipore; cat# 04-820), and GAPDH (0411) (Santa Cruz; cat# sc-47724). RPPA assays were carried out as described previously with minor modifications^32^.

### Statistical Analysis

ANOVA or Student *t* test was used for independent samples with normal distribution. Where distribution was not normal (assessed using Q-Q plots with the Wilk-Shapiro test of normality), either the Kruskal-Wallis or Wilcoxon Rank Sum test was used. All experiments were conducted in triplicate, and each experiment was duplicated independently >2 times. These criteria were formulated to ensure that results from each dataset were calculable within the range of sensitivity of the statistical test used. Databases used for human data mining are from publically available resources: Oncomine, cBio^33^, and COSMIC. Z1031 dataset was used with permission from the Alliance consortium. All patients provided informed consent, and studies were conducted according to ethical guidelines and with Institutional Review Board approval.

MutL^-^ tumor from METABRIC, TCGA, and Z1031 datasets was determined in a case list containing all ER^+^ sample IDs based on gene expression less than mean-1.5 × standard deviation and/or the presence of nonsilent mutations in *MLH1* and *PMS2.* For the multivariate analysis, we analyzed ER^+^ tumor samples, extracting mutation data from the cBio portal, and corresponding clinical data through Oncomine. Only samples with survival metadata were included in the analysis. Gene expression, and survival data for TCGA samples were downloaded from cBio portal. All survival data were analyzed using Kaplan-Meier curves and log-rank tests.

Proportional hazards were determined using Cox regression. Sample size for animal experiments was estimated using power calculations in R. *P* values were adjusted for multiple comparisons where appropriate using Benjamini-Hochberg. All graphs and statistical analyses were generated either in MS Excel or R and edited in Adobe Photoshop or Illustrator. Z1031ClinicalTrials.gov Identifier: NCT00265759. Data for Z1031 samples available in dbGaP (phs000472.v2.p1).

## Acknowledgments

We would like to acknowledge the Patient-derived Xenograft and Advanced *In Vivo* Models core (funded by P30 Cancer Center Support Grant NCI-CA125123, CPRIT Core Facilities Support Grant RP170691) and Dr. Michael T. Lewis, Ph.D., Academic Director, Lacey E. Dobrolecki, MS, Core Director at Baylor College of Medicine for helping us in engrafting WHIM20 PDX explants. We also thank Dr. Alejandra Bruna (CRUK, UK) and Dr. Violeta Serra (VHIO, Barcelona) for providing PDX drug response data and tumor sections for the STG and VHIO PDX lines.

## Funding

Work in this study was funded by Department of Defense Breast Cancer Research Program Breakthrough awards (W81XWH-18-1-0034 to SH, W81XWH-18-1-0040, W81XWH-18-1-0035 and W81XWH-18-1-0084 to SMK), NCI K22 Career Development award (CA229613 to SH), Susan G. Komen Promise grant (PG12220321 to MJE), and Cancer Prevention and Research Institute of Texas (CPRIT) Recruitment of Established Investigators award (RR140033 to MJE), National Cancer Institute of the National Institutes of Health under Award Numbers U10CA180821 and U10CA180882 (to the Alliance for Clinical Trials in Oncology), U24C196171.

## Author contributions

NP designed and performed experiments, analyzed data and helped write the manuscript. SS helped design, conduct and analyze data from Western blots and xenograft experiments. VD helped design and conduct 3D Matrigel assays and immunofluorescence experiments. LS, TP, RK and CC conducted WHIM20 patient-derived xenograft experiment. AM helped conduct immunofluorescence experiments. MJE and SK helped design experiments, interpret results, and edit the manuscript. SH designed and performed experiments, analyzed and interpreted data, and wrote and edited the manuscript.

## Competing interests

MJE has intellectual property ownership and received royalties for the PAM50-based breast cancer test “Prosigna”. In the last 5 years he has received ad hoc consulting fees and meals (less than $5000 per year) from Abbvie, Novartis, AstraZenica, Pfizer, Sermonix, and Puma. MJE is a McNair Foundation Fellow and Susan G. Komen Scholar. SMK is a stakeholder in NeoZenome Therapeutics Inc.

## Data and materials availability

Z1031ClinicalTrials.gov Identifier: NCT00265759. Data for Z1031 samples available in dbGaP (phs000472.v2.p1).

## References and Notes

1. Schiff, R. et al. Cross-talk between estrogen receptor and growth factor pathways as a molecular target for overcoming endocrine resistance. Clin. Cancer Res. Off. J. Am. Assoc. Cancer Res. 10, 331S–6S (2004).

2. Fu, X., De Angelis, C., Veeraraghavan, J., Osborne, C. K. & Schiff, R. Molecular Mechanisms of Endocrine Resistance. in Estrogen Receptor and Breast Cancer: Celebrating the 60th Anniversary of the Discovery of ER (ed. Zhang, X.) 265–307 (Springer International Publishing, 2019). doi:10.1007/978-3-319-99350-8_11.

3. Finn, R. S. et al. Estrogen receptor, progesterone receptor, human epidermal growth factor receptor 2 (HER2), and epidermal growth factor receptor expression and benefit from lapatinib in a randomized trial of paclitaxel with lapatinib or placebo as first-line treatment in HER2-negative or unknown metastatic breast cancer. J. Clin. Oncol. Off. J. Am. Soc. Clin. Oncol. 27, 3908–3915 (2009).

4. Burstein, H. J. et al. Endocrine therapy with or without inhibition of epidermal growth factor receptor and human epidermal growth factor receptor 2: a randomized, double-blind, placebo-controlled phase III trial of fulvestrant with or without lapatinib for postmenopausal women with hormone receptor-positive advanced breast cancer-CALGB 40302 (Alliance). J. Clin. Oncol. Off. J. Am. Soc. Clin. Oncol. 32, 3959–3966 (2014).

5. Ellard, S. L. et al. Randomized phase II study comparing two schedules of everolimus in patients with recurrent/metastatic breast cancer: NCIC Clinical Trials Group IND.163. J. Clin. Oncol. Off. J. Am. Soc. Clin. Oncol. 27, 4536–4541 (2009).

6. Ma, C. X. et al. A Phase II Trial of Neoadjuvant MK-2206, an AKT Inhibitor, with Anastrozole in Clinical Stage II or III PIK3CA-Mutant ER-Positive and HER2-Negative Breast Cancer. Clin. Cancer Res. Off. J. Am. Assoc. Cancer Res. 23, 6823–6832 (2017).

7. ACS Facts and Figures 2017-2018. https://www.cancer.org/content/dam/cancer-org/research/cancer-facts-and-statistics/breast-cancer-facts-and-figures/breast-cancer-facts-and-figures-2017-2018.pdf.

8. Ellis, M. J. Lessons in precision oncology from neoadjuvant endocrine therapy trials in ER+ breast cancer. Breast Edinb. Scotl. (2017) doi:10.1016/j.breast.2017.06.039.

9. Slamon, D. J. et al. Use of Chemotherapy plus a Monoclonal Antibody against HER2 for Metastatic Breast Cancer That Overexpresses HER2. N. Engl. J. Med. 344, 783–792 (2001).

10. Ellis, M. J. & Perou, C. M. The Genomic Landscape of Breast Cancer as a Therapeutic Roadmap. Cancer Discov. 3, 27 (2013).

11. Bose, R. et al. Activating HER2 mutations in HER2 gene amplification negative breast cancer. Cancer Discov. 3, 224–237 (2013).

12. Beaver, J. A. et al. FDA Approval: Palbociclib for the Treatment of Postmenopausal Patients with Estrogen Receptor-Positive, HER2-Negative Metastatic Breast Cancer. Clin. Cancer Res. Off J. Am. Assoc. Cancer Res. (2015) doi:10.1158/1078-0432.CCR-15-1185.

13. Anurag, M., Ellis, M. J. & Haricharan, S. DNA damage repair defects as a new class of endocrine treatment resistance driver. Oncotarget 9, 36252–36253 (2018).

14. Portman, N. et al. Overcoming CDK4/6 inhibitor resistance in ER-positive breast cancer. Endocr. Relat. Cancer 26, R15–R30 (2019).

15. Haricharan, S. et al. Loss of MutL disrupts Chk2-dependent cell cycle control through CDK4/6 to promote intrinsic endocrine therapy resistance in primary breast cancer. Cancer Discov. 7, 1168–1183 (2017).

16. Anurag, M. et al. Comprehensive Profiling of DNA Repair Defects in Breast Cancer Identifies a Novel Class of Endocrine Therapy Resistance Drivers. Clin. Cancer Res. Off. J. Am. Assoc. Cancer Res. 24, 4887–4899 (2018).

17. Brown, T. C. & Jiricny, J. Repair of base-base mismatches in simian and human cells. Genome 31, 578–583 (1989).

18. Wardell, S. E. et al. Efficacy of SERD/SERM Hybrid-CDK4/6 Inhibitor Combinations in Models of Endocrine Therapy-Resistant Breast Cancer. Clin. Cancer Res. Off. J. Am. Assoc. Cancer Res. 21, 5121–5130 (2015).

19. Haricharan, S., Bainbridge, M. N., Scheet, P. & Brown, P. H. Somatic mutation load of estrogen receptor-positive breast tumors predicts overall survival: an analysis of genome sequence data. Breast Cancer Res. Treat. 146, 211–220 (2014).

20. Bruna, A. et al. A Biobank of Breast Cancer Explants with Preserved Intra-tumor Heterogeneity to Screen Anticancer Compounds. Cell 167, 260–274.e22 (2016).

21. Ellis, M. J. et al. Randomized phase II neoadjuvant comparison between letrozole, anastrozole, and exemestane for postmenopausal women with estrogen receptor-rich stage 2 to 3 breast cancer: clinical and biomarker outcomes and predictive value of the baseline PAM50-based intrinsic subtype--ACOSOG Z1031. J. Clin. Oncol. Off. J. Am. Soc. Clin. Oncol. 29, 2342–2349 (2011).

22. Vera-Ramirez, L. et al. Oxidative stress status in metastatic breast cancer patients receiving palliative chemotherapy and its impact on survival rates. Free Radic. Res. 46, 2–10 (2012).

23. Finn, R. S. et al. Quantitative ER and PgR Assessment as Predictors of Benefit from Lapatinib in Postmenopausal Women with Hormone Receptor-Positive, HER2-Negative Metastatic Breast Cancer. Clin. Cancer Res. 20, 736–743 (2014).

24. Mismatch Repair Protein Loss as a Prognostic and Predictive Biomarker in Breast Cancers Regardless of Microsatellite Instability. - PubMed - NCBI. https://www.ncbi.nlm.nih.gov/pubmed/31360876.

25. Lanza, G. et al. Immunohistochemical pattern of MLH1/MSH2 expression is related to clinical and pathological features in colorectal adenocarcinomas with microsatellite instability. Mod. Pathol. Off. J. U. S. Can. Acad. Pathol. Inc 15, 741–749 (2002).

26. Stelloo, E. et al. Practical guidance for mismatch repair-deficiency testing in endometrial cancer. Ann. Oncol. Off. J. Eur. Soc. Med. Oncol. 28, 96–102 (2017).

27. Kloth, M. et al. Activating ERBB2/HER2 mutations indicate susceptibility to pan-HER inhibitors in Lynch and Lynch-like colorectal cancer. Gut 65, 1296–1305 (2016).

28. Win, A. K. et al. Risks of colorectal and other cancers after endometrial cancer for women with Lynch syndrome. J. Natl. Cancer Inst. 105, 274–279 (2013).

29. Ghanipour, L., Jirström, K., Sundström, M., Glimelius, B. & Birgisson, H. Associations of defect mismatch repair genes with prognosis and heredity in sporadic colorectal cancer. Eur. J. Surg. Oncol. J. Eur. Soc. Surg. Oncol. Br. Assoc. Surg. Oncol. 43, 311–321 (2017).

30. Haricharan, S. & Brown, P. TLR4 has a TP53-dependent dual role in regulating breast cancer cell growth. Proc. Natl. Acad. Sci. U. S. A. 112, E3216–3225 (2015).

31. Haricharan, S. et al. Mechanism and preclinical prevention of increased breast cancer risk caused by pregnancy. eLife 2, e00996 (2013).

32. Chang, C.-H. et al. Mammary Stem Cells and Tumor-Initiating Cells Are More Resistant to Apoptosis and Exhibit Increased DNA Repair Activity in Response to DNA Damage. Stem Cell Rep. 5, 378–391 (2015).

33. Cerami, E. et al. The cBio Cancer Genomics Portal: An Open Platform for Exploring Multidimensional Cancer Genomics Data. Cancer Discov. 2, 401–404 (2012).

