## Supplementary material for "Mismatch repair deficiency predicts response to HER2 blockade in HER2-negative breast cancer": Supp Fig 1

### Supplementary Materials:

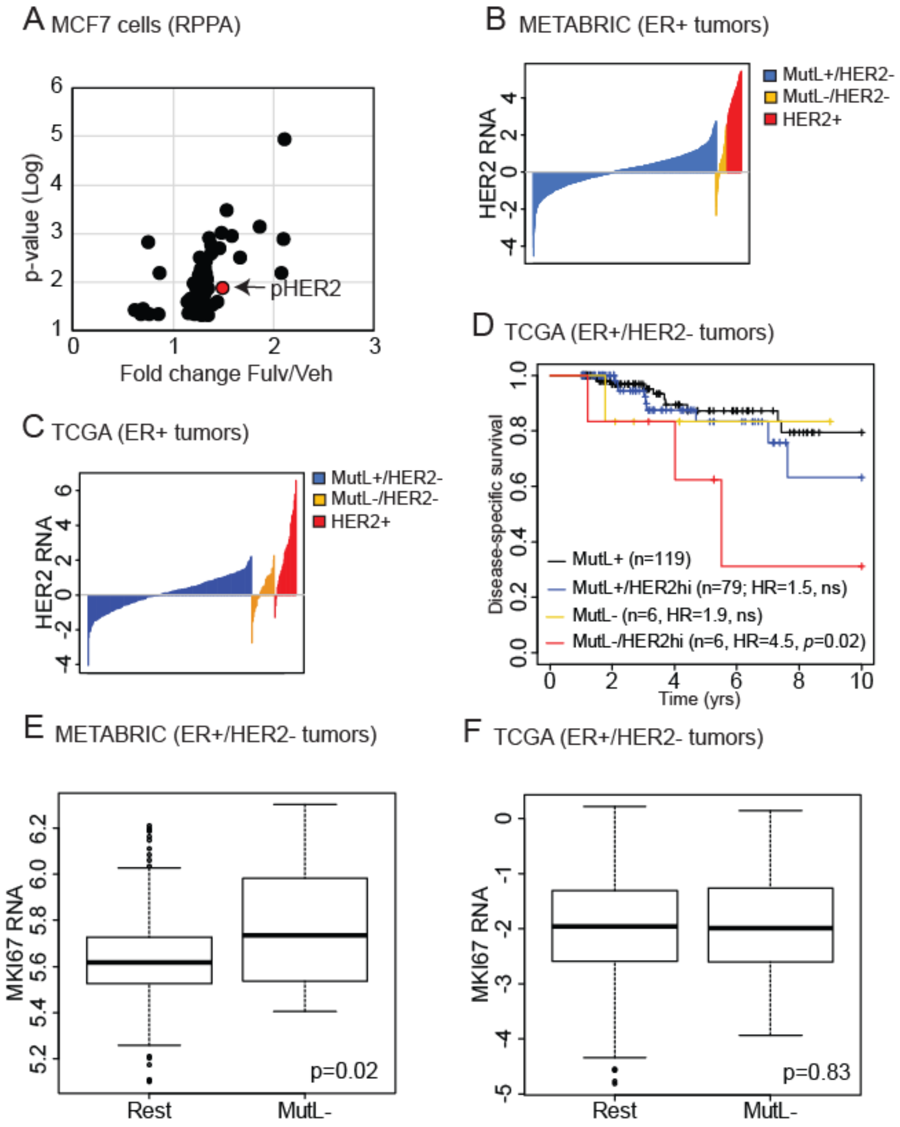


**Supplementary Figure 1:** $ER^+$, nominally $HER2^-$ breast cancer patients whose tumors are $MutL^-$ have relatively high levels of HER2 and significantly worse disease-specific outcomes. (A) RPPA data of MCF7 sh*Luc*, sh*MLH1* and sh*PMS2* cells grown +/- fulvestrant and analyzed for proteins whose levels increase specifically in response to fulvestrant in sh*MutL* cells relative to sh*Luc*. P-value generated using Student's t-test and corrected for multiple comparison using Bonferroni. Four replicates assayed. (B-C) Index plot depicting RNA levels of *HER2* in $MutL^-$ (gold) and $MutL^+$ (blue) $ER^+$/nominally $HER2^-$ patient tumors from METABRIC (B) and TCGA (C). $HER2^+$ (or amplified) patient tumors (red) are included to provide context. Supports data in Fig 1A. (D) Kaplan-Meier survival curves of indicated groups of patients from TCGA demonstrating differences in disease-specific survival. Cox Regression analysis determined p-values and hazard ratios. Supports data in Fig 1B. (E-F) Boxplots demonstrating comparable levels of gene expression of the proliferation marker, Ki67 in two independent datasets between MutL- and $MutL^+$ $ER^+$/HER2-non-amplified patient tumors. Wilcoxon Rank sum test determined p-values. Horizonal line represents the mean and error bars the standard error.
