## Supplementary material for "Mismatch repair deficiency predicts response to HER2 blockade in HER2-negative breast cancer": Supp Fig 2

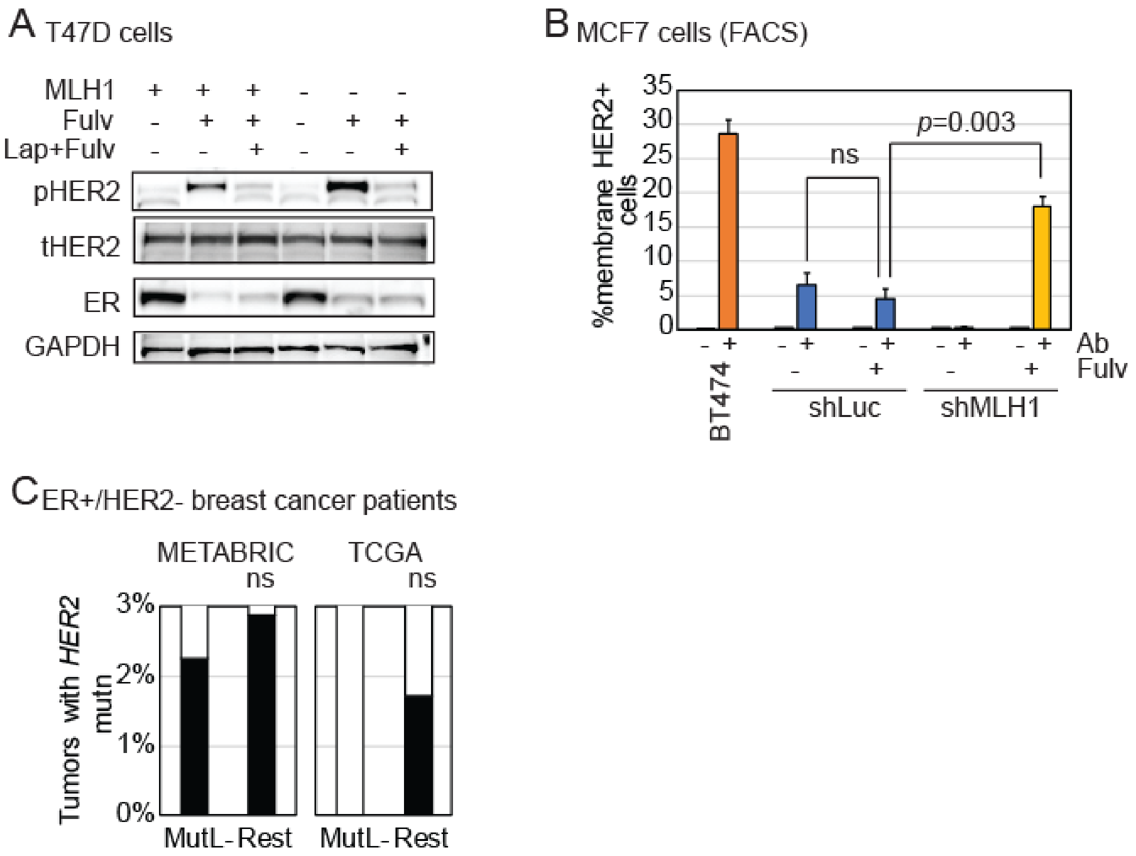


**Supplementary Figure 2:** MLH1 loss in $ER^+$, nominally $HER2^-$ breast cancer cells causally induces membrane-bound HER2 upon endocrine treatment. (A) Western blot demonstrating increased pHER2 levels in sh*Luc* and sh*MLH1* T47D cells treated with vehicle, fulvestrant (Fulv) or a combination of lapatinib, a HER inhibitor, and fulvestrant (Lap+Fulv). Supports data in Fig 2A. (B) Bar graph representing percent cells with membrane bound HER2 detected by membrane FACS in sh*Luc* and sh*MLH1* MCF7 cells treated with or without fulvestrant (Fulv). Each group has an isotype control (Ab) and BT474, HER2-amplied cells are included as positive control. Columns represent the mean and error bars the standard deviation. Student's t-test determined p-values. Supports data in Fig 2B. (C) Incidence of HER2 mutations in $ER^+$/HER2-non-amplified breast tumors from two independent patient cohorts. Fisher's exact test determined p-values.
