## Supplementary material for "Mismatch repair deficiency predicts response to HER2 blockade in HER2-negative breast cancer": Supp Fig 3

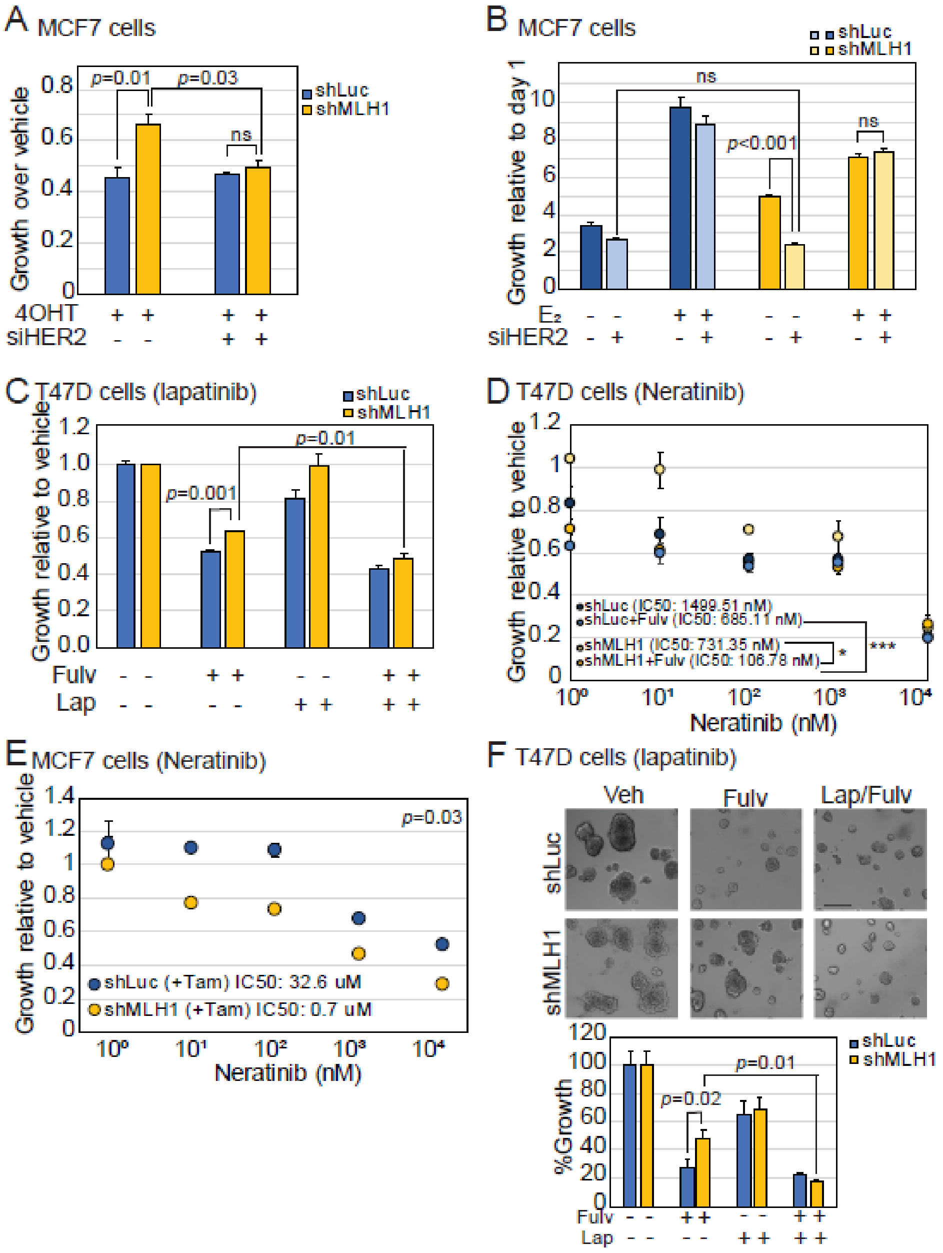


**Supplementary Figure 3:** HER2 is required for endocrine treatment resistant growth of ER$^+$ MLH1$^-$ breast cancer cells. (A-B) Bar graphs representing relative growth of MCF7 sh*Luc* and sh*MLH1* cells transiently transfected with either scrambled siRNA or siRNA against *HER2*, and then treated with tamoxifen (4-OHT, A) or grown in charcoal stripped serum and deprived of estrogen ($E_2$, B). Western blotting validating knockdown and response to fulvestrant presented in Fig 3A-B. (C) Bar graphs demonstrating increased sensitivity of T47D sh*MLH1* cells to a combination of fulvestrant (Fulv) and lapatinib (Lap). Supports data presented in Fig 3C. (D-E) Dose response curves of sh*Luc* and sh*MLH1* T47D (D) and MCF7 (E) cells treated with neratinib in combination with fulvestrant (Fulv, D) or tamoxifen (Tam, E). IC50 values were calculated and differences in IC50 from three independent experiments was statistically compared. Circles represent mean relative growth and error bars the standard deviation. Supports data presented in Fig 3D. (F) 3D growth of sh*Luc* and sh*MLH1* T47D cells in Matrigel shown with representative photomicrographs and accompanying quantification. Scale bars represent 50μ. Supports data presented in Fig 3E. For all bar graphs, bars represent the mean and error bars the standard deviation. Student's t-test determined all p-values.
