## Supplementary material for "Mismatch repair deficiency predicts response to HER2 blockade in HER2-negative breast cancer": Supp Fig 4

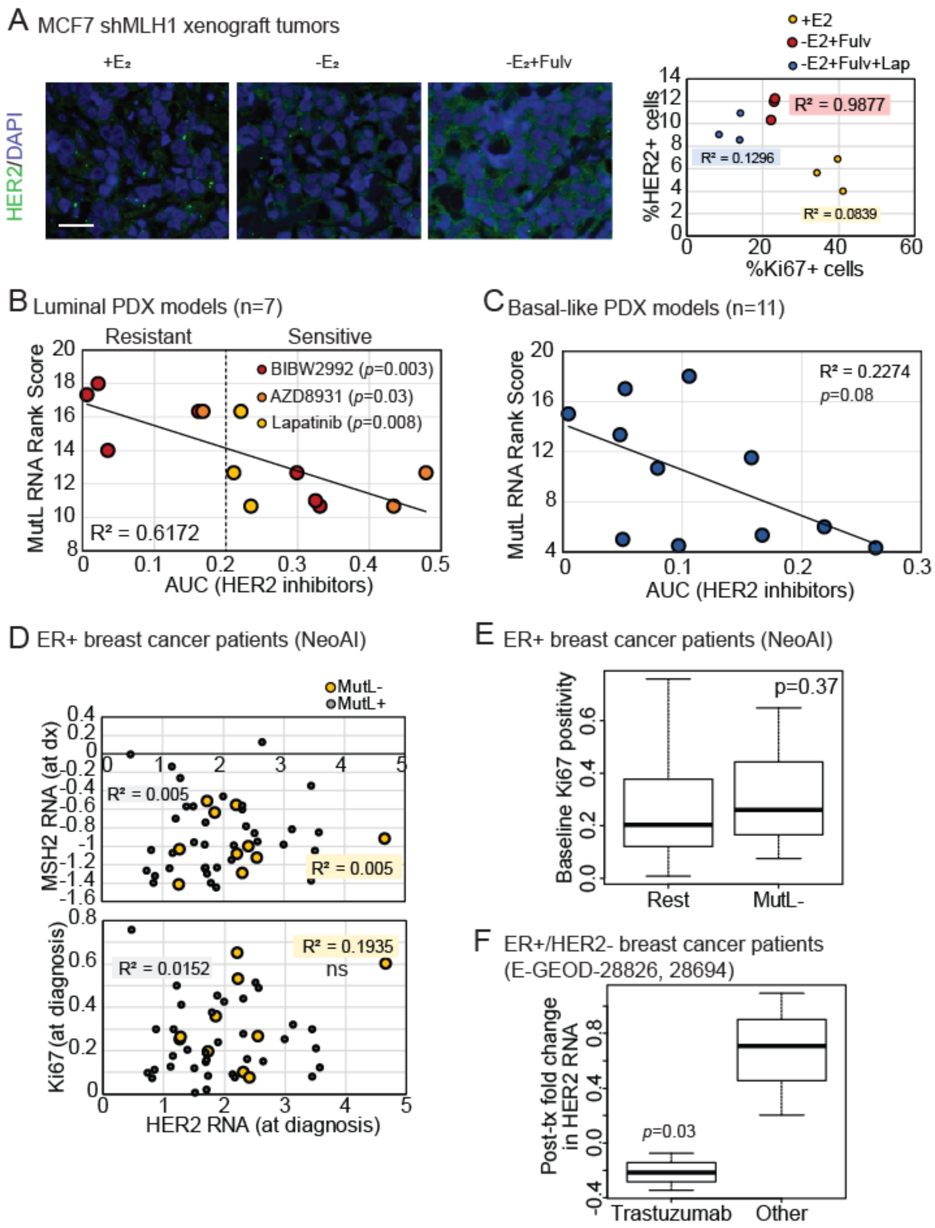


**Supplementary Figure 4:** MLH1 loss predicts sensitivity to HER2 inhibitors in endocrine treatment resistant $ER^+$, nominally $HER2^-$ breast cancer cells *in vivo* and in patient tumors. (A) Immunofluorescent analysis of HER2 in MCF7 sh*MLH1* xenograft tumors administered with specified therapies depicted with representative photomicrographs. Scale bar represents 50µ. Accompanying regression analysis of correlation between HER2 positivity and levels of Ki67, a proliferation marker. Supports data in Fig 4A. (B-C) Regression analysis depicting lack of significant correlation between *MLH1* and *PMS2* RNA levels, and sensitivity to HER2 inhibitors in luminal (B) and basal-like (C) PDX tumors grown *in vivo*. Supports data in Fig 4B. (D) Regression analysis of correlation between *MSH2* RNA levels (top) and proliferation marker, Ki67 before endocrine treatment (bottom) with *HER2* RNA levels. Supports data in Fig 4C. For all graphs, linear regression model analysis was used to determine $R^2$ and p-value. (E-F) Boxplots demonstrating comparable levels of Ki67 positivity between $MutL^-$ and $MutL^+$ $ER^+/HER2^-$ breast tumors from the NeoAI database (E) and downregulation of *HER2* RNA levels in response to Herceptin, but not in response to anthracyclines or taxanes (Other, F). Supports data in Fig 4D. Line represents median and error bars the standard error of the mean. Wilcoxon Rank Sum test determined p-value.
